# Probing multi-way chromatin interaction with hypergraph representation learning

**DOI:** 10.1101/2020.01.22.916171

**Authors:** Ruochi Zhang, Jian Ma

## Abstract

Advances in high-throughput mapping of 3D genome organization have enabled genome-wide characterization of chromatin interactions. However, proximity ligation based mapping approaches for pairwise chromatin interaction such as Hi-C cannot capture multi-way interactions, which are informative to delineate higher-order genome organization and gene regulation mechanisms at single-nucleus resolution. The very recent development of ligation-free chromatin interaction mapping methods such as SPRITE and ChIA-Drop has offered new opportunities to uncover simultaneous interactions involving multiple genomic loci within the same nuclei. Unfortunately, methods for analyzing multi-way chromatin interaction data are significantly underexplored. Here we develop a new computational method, called MATCHA, based on hypergraph representation learning where multi-way chromatin interactions are represented as hyperedges. Applications to SPRITE and ChIA-Drop data suggest that MATCHA is effective to denoise the data and make *de novo* predictions of multi-way chromatin interactions, reducing the potential false positives and false negatives from the original data. We also show that MATCHA is able to distinguish between multi-way interaction in a single nucleus and combination of pairwise interactions in a cell population. In addition, the embeddings from MATCHA reflect 3D genome spatial localization and function. MATCHA provides a promising framework to significantly improve the analysis of multi-way chromatin interaction data and has the potential to offer unique insights into higher-order chromosome organization and function.

## Introduction

Interphase chromosomes in higher eukaryotic cells are folded and packaged in the nucleus, leading to higher-order chromatin interactions in three-dimensional (3D) space that are key to understanding gene regulation and cell function (Kumaran et al., 2008; Bonev and Cavalli, 2016). Advances in high-throughput mapping of nuclear genome organization by capturing pairwise interactions between proximal genomic loci, e.g., Hi-C (Lieberman-Aiden et al., 2009; Rao et al., 2014) and ChIA-PET (Fullwood and Ruan, 2009; Tang et al., 2015), have enabled genome-wide characterization of 3D genome features, including loops (Rao et al., 2014; Tang et al., 2015), topologically associating domains (Dixon et al., 2012; Nora et al., 2012), A/B compartments (Lieberman-Aiden et al., 2009) and subcompartments (Rao et al., 2014; Xiong and Ma, 2019). However, one limitation of the proximity ligation based methods is that they capture interactions between genomic loci that are in close proximity to directly ligate and are unable to reveal chromatin interactions that are beyond the distance of direct ligation (Beagrie et al., 2017; Quinodoz et al., 2018). More importantly, most widely available Hi-C and ChIA-PET data only measure pairwise interactions and cannot delineate multiple chromatin loci that interact simultaneously in the same nucleus (Kempfer and Pombo, 2019).

Very recently, new technologies such as SPRITE (Quinodoz et al., 2018) and ChIA-Drop (Zheng et al., 2019) have been developed to capture simultaneous interactions among multiple genomic loci within individual nuclei. Based on the SPRITE data, Quinodoz et al. (2018) reported that the inter-chromosomal chromatin interactions can be partitioned into distinct active and inactive hubs. ChIA-Drop, on the other hand, allows the detection of multi-way chromatin interactions mediated by specific proteins (Zheng et al., 2019). For example, from RNA PolII ChIA-Drop data, potential co-regulated genes can be characterized. These recently developed methods have demonstrated the unique properties of the 3D genome architecture that can only be manifested by multi-way chromatin interactions (Beagrie et al., 2017; Quinodoz et al., 2018; Zheng et al., 2019).

However, there are a number of unsolved challenges in analyzing multi-way chromatin interaction data. First, existing methods for analyzing SPRITE and ChIA-Drop data typically have strong assumptions that would lead to loss of information from the original data. For example, Peakachu (Salameh et al., 2019) extracts chromatin interactions from multi-way interaction data by simply decomposing clusters into a pairwise contact matrix to directly apply methods developed for Hi-C and ChIA-PET, which causes a dramatic loss of higher-order contact information. Recently, MIA-Sig (Kim et al., 2019) was developed to remove noise and call significant chromatin complex in ChIA-Drop data based on the assumption that the genomic distance of a true multi-way interaction should be more evenly spaced and if a genomic locus is far from other loci within the droplet then it is likely contamination. This assumption, however, may introduce errors and has yet to be further evaluated and confirmed (Kim et al., 2019). Second, the observed frequencies for clusters with a larger number of loci are much lower than smaller ones, making it increasingly more difficult to reliably denoise the data based on frequencies only. For example, in Quinodoz et al. (2018), combinations of 1Mb genomic bins (referred to as *k*-mers) were required to be observed at least in 5 SPRITE clusters (occurrence frequency ≥ 5) to be further considered. However, when analyzing multi-way interactions with larger size and higher resolution, the cut-off for frequencies of larger *k*-mers would be much harder to determine because of their low occurrence. Third, the number of multi-way interaction datasets and their quality remain limited, but no reliable predictive method is currently available to either denoise the data or to make *de novo* multi-way interaction predictions. Indeed, in order to fully realize the potential of the emerging multi-way chromatin interaction data, it is imperative to have new algorithms that can extract important patterns by addressing the aforementioned challenges.

Here we develop a new generic computational framework, called MATCHA (Multi-wAy inTeracting CHromatin Analysis), for the analysis of multi-way chromatin interaction data. We consider the 3D genome as a graph where nodes are genomic bins and edges connecting bins represent chromatin interactions between bins. When an interaction involves more than two nodes, it is referred to as a hyperedge and the graph containing hyperedges is called a hypergraph (Berge, 1984; Zhou et al., 2007). In other words, we model each multi-way interaction as a hyperedge. MATCHA takes multi-way chromatin interaction data as input and extracts patterns from the corresponding hypergraph. The patterns are represented as embedding vectors for each genomic bin that reflect the properties of 3D chromatin structures. The model can further predict the probability for a group of genomic bins having simultaneous interactions, for either denoising the dataset or making *de novo* predictions. Taken together, MATCHA is a new computational method based on hypergraph representation learning for the analysis of multi-way chromatin interaction data that can provide new insights into nuclear genome structure and function. Source code of MATCHA can be accessed at: https://github.com/ma-compbio/MATCHA.

## Results

### Overview of the MATCHA algorithm

Fig. 1A illustrates the workflow of MATCHA for the analysis of multi-way chromatin interaction data. There are four main components: (1) Constructing hypergraphs (the formal definition can be found in the Methods section) based on the multi-way chromatin interaction data where non-overlapping genomic bins are defined as nodes and bins in the same multi-way interaction are connected by hyperedges. (2) Generating node features for the hypergraph based on decomposed pairwise contact matrix from multiway interaction data. The decomposed pairwise contact matrix then passes through the Mix-n-Match autoencoder (see Methods). (3) Generating labeled data for the training of the hypergraph representation learning model. Within the dataset, positive samples are defined as existing hyperedges while negative samples are unobserved ones. We generate negative samples through an efficient and biologically meaningful negative sampling strategy. (4) Training our hypergraph representation learning model Hyper-SAGNN (Zhang et al., 2020) which takes both labeled data and node features as input (Fig. 1B). Details of each component are described in the Methods section.

**Figure 1:**
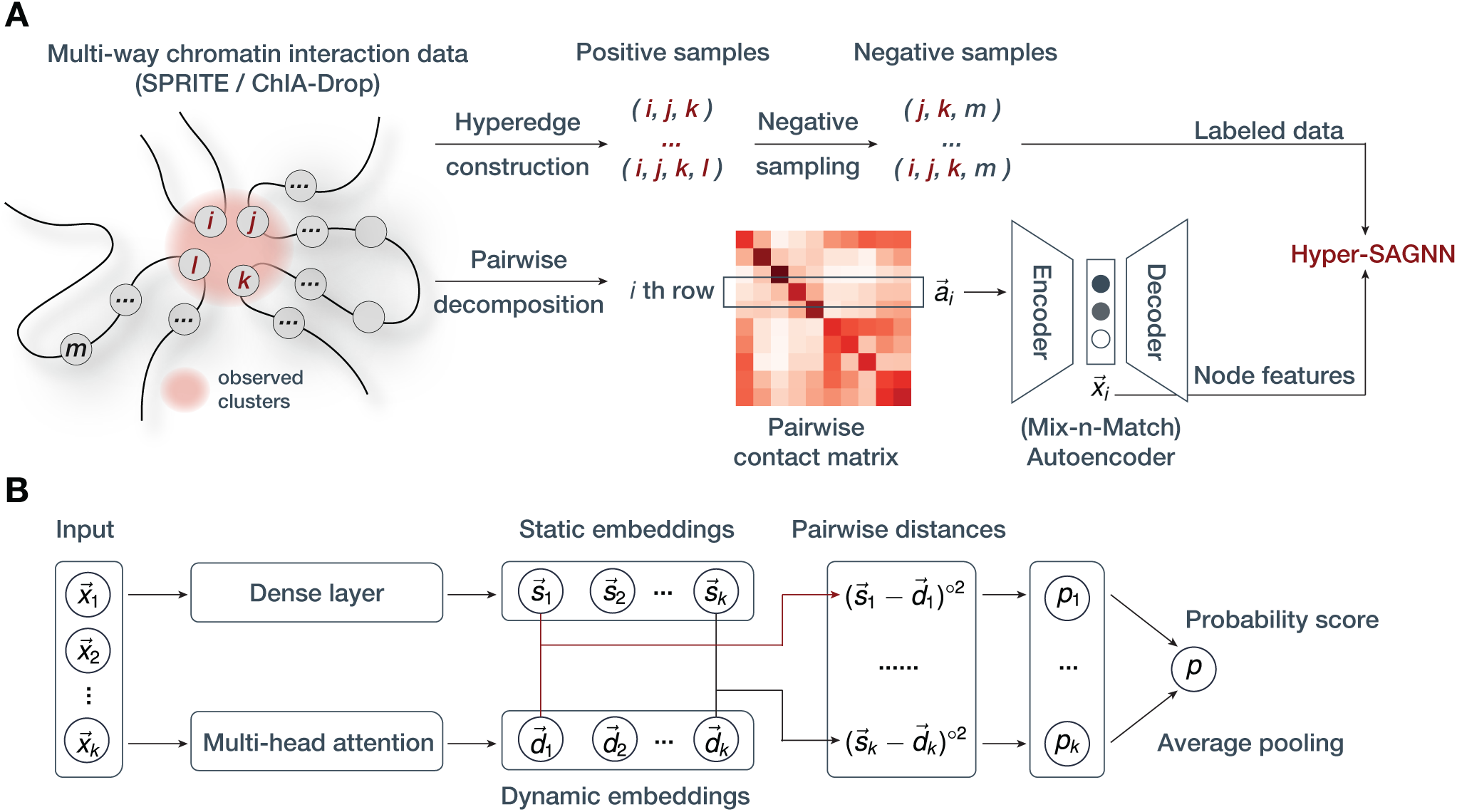
**(A)** Overview of MATCHA. Observed clusters from multi-way chromatin interaction data such as SPRITE and ChIA-Drop are defined as positive samples (hyperedges) and the unobserved groups of nodes are sampled as negative ones. Observed clusters are decomposed into a pairwise contact matrix to generate node features by going through an autoencoder. Both labeled data and node features are used to train Hyper-SAGNN (Zhang et al., 2020) to model the hypergraph constructed by multi-way chromatin interaction data. **(B)** Hyper-SAGNN architecture (details in Zhang et al. (2020)). The input of the model contains groups of nodes with node features, 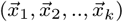. The tuple passes through fully-connected layers and multi-head attention layers to generate static and dynamic node embeddings, respectively. Then a pseudo euclidean distance is calculated for each pair of static/dynamic embeddings which would be turned into probability scores between 0 to 1. The final probability score indicating whether this group of nodes (1, 2, …, *k*) would form a hyperedge is calculated by average pooling.

We remark that in the last step we use our recently developed model, Hyper-SAGNN (Zhang et al., 2020), instead of other previously developed graph representation learning methods for several reasons. The hyperedge prediction problem (the formal definition can be found in the Methods section) is equivalent to learning the function *p* that takes tuples of node features (*x*_1_, …, *x*_*k*_) as input and produces the probability of these nodes forming a hyperedge. Since the number of genomic bins involved in a multi-way interaction varies, the constructed hypergraph would be a non-*k*-uniform hypergraph. Moreover, there is no intrinsic order in each hyperedge; in other words, the function *p* should satisfy *p*(*x*_1_, …, *x*_*k*_) = *p*(shuffle(*x*_1_, …, *x*_*k*_)) where shuffle() represents random shuffling of the order of the nodes involved in the hyperedge. None of the graph representation learning methods for pairwise interactions (such as DeepWalk (Perozzi et al., 2014) and node2vec (Grover and Leskovec, 2016)) or previously developed hyperedge prediction methods (such as DHNE (Tu et al., 2018) and HEBE (Gui et al., 2016)) can handle our situation because they either cannot model higher-order information within hyperedges or require fixed-sized and ordered input. We recently developed Hyper-SAGNN (Zhang et al., 2020) for the general hypergraph representation learning problem and it is applicable to homogeneous and heterogeneous hypergraphs with variable hyperedge sizes. We demonstrated in Zhang et al. (2020) that Hyper-SAGNN achieves state-of-the-art performance in multiple applications.

### MATCHA accurately predicts multi-way chromatin interactions with different sizes

We first briefly describe the data we used in this work and the construction of hypergraphs. The SPRITE data are from the GM12878 lymphoblastoid human cell line Quinodoz et al. (2018). The RNAPII enriched ChIA-Drop data are from *Drosophila* S2 cells (Zheng et al., 2019). Unless otherwise stated, we used 1Mb resolution for the SPRITE data and 5kb resolution for the RNAPII ChIA-Drop data to build the hypergraph. The details of data processing are described in the Methods section. In Table 1, we list the number of different sized hyperedges for both datasets grouped by the occurrence frequency. As expected, the number of hyperedges decreases with the larger size of hyperedge and larger occurrence frequency cut-off. To balance the number of hyperedges over different sizes in SPRITE data, we selected hyperedges that have occurrence frequency ≥ 8 for hyperedges with the size of 3, ≥ 3 for hyperedges with the size of 4, and ≥ 2 for hyperedges with the size of 5. This resulted in around 600k hyperedges of size 3, 700k hyperedges of size 4, and 500k hyperedges of size 5. Note that using a more stringent standard to define hyperedges would lead to higher accuracy, but this also leads to smaller training samples and potential bias because hyperedges with larger occurrence frequency tend to have smaller genomic distance. For the RNAPII ChIA-Drop data, all hyperedges are “intra” as the inter-chromosomal reads were filtered in the original ChIA-Drop pipeline. As mentioned in the Methods section and shown in Table 1, the remaining number of hyperedges decreases more dramatically for RNAPII ChIA-Drop data as compared to SPRITE. We therefore used 2 as the cut-off for hyperedges of all sizes.

**Table 1:** Overview of the constructed hypergraphs from SPRITE data and RNAPII ChIA-Drop data. The number of hyperedges for different size over different occurrence frequency group are reported in the table.

| SPRITE |  |  |  | RNAPII ChIA-Drop |  |  |  |
| --- | --- | --- | --- | --- | --- | --- | --- |
| Occ_freq | 3 | 4 | 5 | Occ_freq | 3 | 4 | 5 |
| 2 | 61,997,214 | 3,823,725 | 507,375 | 1 | 1,022,258 | 1,928,187 | 4,321,741 |
| 3-4 | 7,953,184 | 591,427 | 17,532 | 2 | 29,145 | 8,603 | 2,631 |
| 5-7 | 795,940 | 89,938 | 1,705 | > 2 | 6,491 | 528 | 32 |
| 8-12 | 342,242 | 12,871 | 397 |  |  |  |  |
| > 12 | 317,195 | 3,687 | 52 |  |  |  |  |

We evaluated MATCHA based on the hypergraph constructed from the SPRITE data with 80% of the hyperedges as the training samples and 20% of them as the testing samples. We quantified the performance by AUROC (area under the receiver operating characteristic) and AUPR (area under the precision-recall curve) scores on the testing dataset. Both metrics were calculated for hyperedges with different sizes separately to provide a comprehensive assessment. The performance is shown in Table 2. We found that MATCHA is able to make accurate predictions across hyperedges with different sizes. We observed lower AUROC and AUPR score for hyperedges of size 5, which may be caused by the lower occurrence frequency cut-off for larger hyperedges. We also performed the evaluation based on the hypergraph constructed from RNAPII ChIA-Drop data. Our method again achieves strong performance in terms of the prediction of hyperedges. However, we observed a different trend of performance versus the size of hyperedge compared with the results from SPRITE, which could be due to the uniform cut-off we used for RNAPII ChIA-Drop data whereas we used a lower cut-off for larger hyperedges for the SPRITE data. Overall, this evaluation suggests that MATCHA is able to accurately predict hyperedges solely based on a fraction of the hyperedges derived from multi-way chromatin interaction data.

**Table 2:** AUROC and AUPR score on the test set for hyperedge prediction. The ratio of positive samples vs. negative samples is 1:3. AUROC and AUPR score are calculated for different size of hyperedges separately to provide a more comprehensive assessment of the performances.

| Data | Group | AUROC | AUPR | Data | Group | AUROC | AUPR |
| --- | --- | --- | --- | --- | --- | --- | --- |
| SPRITE | size = 3 | 0.991 | 0.978 | RNAPII ChIA-Drop | size = 3 | 0.913 | 0.790 |
| SPRITE | size = 4 | 0.949 | 0.869 | RNAPII ChIA-Drop | size = 4 | 0.959 | 0.956 |
| SPRITE | size = 5 | 0.853 | 0.683 | RNAPII ChIA-Drop | size = 5 | 0.977 | 0.987 |

### MATCHA can denoise multi-way chromatin interaction data

Next, we sought to ask if MATCHA can be used to assess whether the clusters with occurrence below the cut-off are real interactions. As a proof-of-principle, we first predict the probability of triplets (3-way interactions) from the occurrence frequency category of 2, 3-4 and 5-7 in Table 1. Note that here we used triplets with occurrence frequency ≥8 as training data. We evaluated the predicted probabilities by comparing with the Hi-C contact matrix. The rationale of this evaluation is that if certain groups of genomic loci simultaneously interact frequently in the cell population, it is expected to see that Hi-C can capture pairwise interactions between each pair of genomic bins.

We first decomposed all the triplets in the training data (triplets with occurrence frequency ≥8) into pairwise edges and identified the corresponding entry in the Hi-C contact matrix. We then calculated the average value of these entries for intra-chromosomal interactions and inter-chromosomal interactions, respectively. These two averaged values were then used to binarize the Hi-C contact matrix and build the Hi-C graph. All the triplets were grouped by the predicted probability score where each group is further categorized by the number of Hi-C edges within the triplets (see Fig. 2A where we also include the positive samples used to train the model as a reference). As shown in Fig. 2A, triplets assigned with higher probability scores are typically enriched with Hi-C edges. For triplets with probability scores *>*0.9, there are more than 40% that have all three pairwise interactions supported by Hi-C edges.

**Figure 2:**
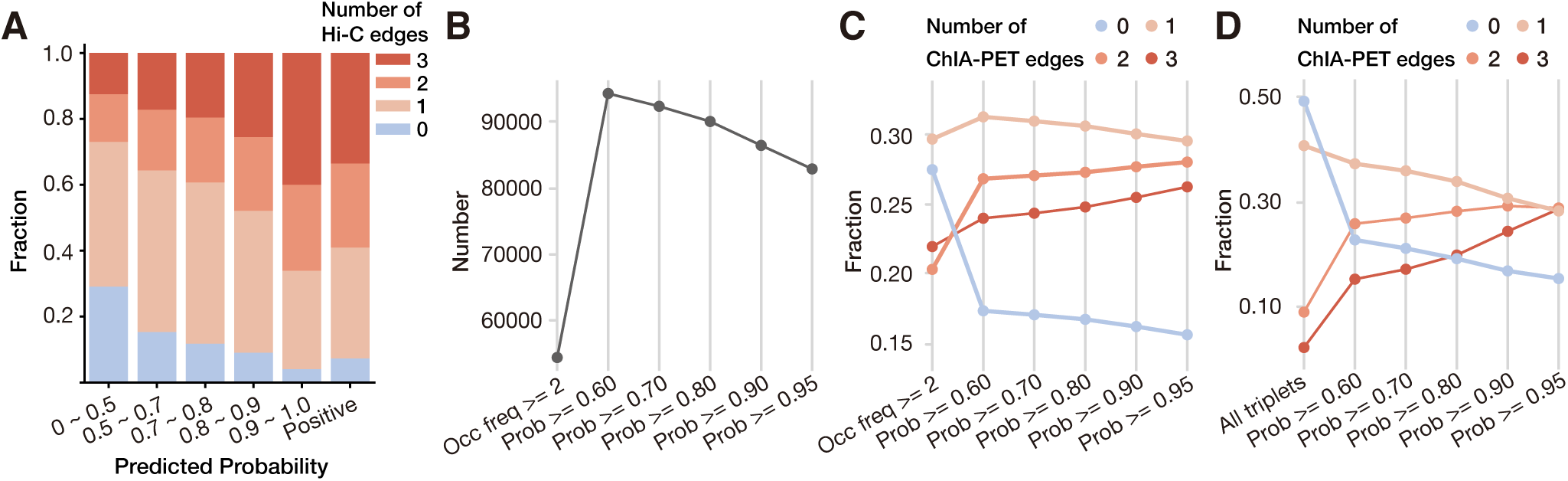
Evaluation of MATCHA’s performence in denoising multi-way chromatin interaction data. **(A)** Distribution of the number of Hi-C edges between the nodes from the triplets using the SPRITE data. The triplets that are observed in SPRITE clusters for frequency between 2-7 are grouped by the predicted probability score. The triplet group that is used in the training as positive samples (observed in SPRITE clusters for more than 8 times) is marked as “positive”. **(B)** The number of triplets in the RNAPII ChIA-Drop data that either have occurrence frequency larger than 2 or have probability score greater than or equal to the listed threshold. **(C)** Distribution of the number of RNAPII ChIA-PET edges between the nodes from the triplets observed in RNAPII ChIA-Drop data. **(D)** Distribution of the number of RNAPII ChIA-PET edges between the nodes from the triplets that are unobserved in RNAPII ChIA-Drop data but have 1D genomic distance within the range 1D genomic distance in the positive triplets.

We then assessed the effectiveness of our method for denoising the RNAPII ChIA-Drop data. Similar to the denoising evaluation on the SPRITE data, we trained the model on hyperedges with occurrence frequency ≥2 as positive samples and then predicted probability scores for all the observed triplets to evaluate the reliability of the predictions by counting the number of *in situ* RNAPII ChIA-PET edges (data from Zheng et al. (2019)) within the triplets. We specifically compared the triplets filtered by the predicted probability score versus the triplets filtered by occurrence frequency. As shown in Fig 2B-C, we found that by using our method for denoising, we can identify more triplets with more ChIA-PET support than simply using occurrence frequency. In addition, although a higher probability cut-off would lead to fewer triplets, it would also result in a higher percentage of triplets that have at least 2 ChIA-PET edges between pairs of genomic bins.

Together, these results demonstrate the great potential of MATCHA as a denoising method. By training the model on a relatively small set of reliable hyperedges, MATCHA can more reliably remove false-positive hyperedges by predicting the probability score for the nodes whose occurrence frequencies are non-zero but not high enough to be confidently assigned as positive samples. As compared to filtering by occurrence frequency, MATCHA has a more principled approach to identify reliable hyperedges.

### MATCHA makes *de novo* predictions of multi-way chromatin interaction

We then asked whether MATCHA is able to make *de novo* predictions, i.e., to predict new hyperedges that are not observed in the original data. Because the hypergraph constructed by RNAPII ChIA-Drop only contains intra-chromosomal hyperedges and has a maximum 1D genomic distance for interactions, it is therefore possible to practically enumerate all the combinations of triplets. We specifically excluded the observed triplets from this triplet set and obtained probability scores based on MATCHA. We again evaluated the reliability of the predictions by comparing to RNAPII ChIA-PET data. In Fig 2D, we found that the triplets with higher probability scores again are more enriched with ChIA-PET edges as compared to all potential triplets. Based on the support from ChIA-PET edges, these triplets with high probability score predicted by MATCHA could potentially be true multi-way interactions that are missed by the original ChIA-Drop data (i.e., false negatives). Note that the fraction of triplets with only 1 ChIA-PET edge is larger than those with 2 or 3 ChIA-PET edges, but this is likely caused by the limited coverage of the RNAPII ChIA-PET data. Specifically, for the data used in this work, there are more than 2M clusters identified from the RNAPII ChIA-Drop data but only around 200K edges from the RNAPII ChIA-PET data. However, the tendency that triplets with higher predicted probability scores would obtain more support from RNAPII ChIA-PET edges demonstrates the potential of MATCHA in making *de novo* predictions from existing data. Based on the model trained from the observed multiway interactions, MATCHA is able to make *de novo* predictions of hyperedges, which could help reveal potential multi-way chromatin interactions undetected in the original data, further showing the advantage of MATCHA as a predictive model compared with other denoising approaches.

### MATCHA improves overall data quality of SPRITE and ChIA-Drop

To further assess the performance of MATCHA for denoising and *de novo* prediction, we used MATCHA to generate a denoised pairwise contact map and compared to either the original contact map or the ones from other data sources with higher coverage. We first evaluated MATCHA on the SPRITE data from GM12878. To achieve a more detailed comparison, we changed the resolution from 1Mb to 100Kb while the other processing procedures remained the same. All pairs of genomic bins were used as input for MATCHA to predict the probabilities of being pairwise interactions, i.e., a “probability map”. We then calculated the element-wise product of the “probability map” and the original contact map, which becomes the denoised contact map. To show the impact of denoising, we compared the original and the denoised SPRITE contact maps to the Hi-C contact map in GM12878. Fig. 3A shows the heatmap comparison of the original and the denoised SPRITE versus Hi-C on chromosome 1, respectively. We found that the denoised SPRITE contact map produced by MATCHA, while preserving similar near-diagonal structures, contains much less noise and clearer patterns for long-range interactions. These off-diagonal patterns based on MATCHA also correspond well with the Hi-C contact map. Detailed quantification of the similarity between the original and the denoised SPRITE versus Hi-C is shown in Fig. 3B, where we utilized the stratum adjusted correlation coefficient (SCC) used in HiCRep (Yang et al., 2017)) with different maximum distance, Pearson correlation score, and Spearman correlation score. The denoised contact map achieves higher scores for all similarity metrics than the original SPRITE contact map compared with the Hi-C contact map. In particular, the advantage is more pronounced with metrics that take into account long-range interactions (Fig. 3B).

**Figure 3:**
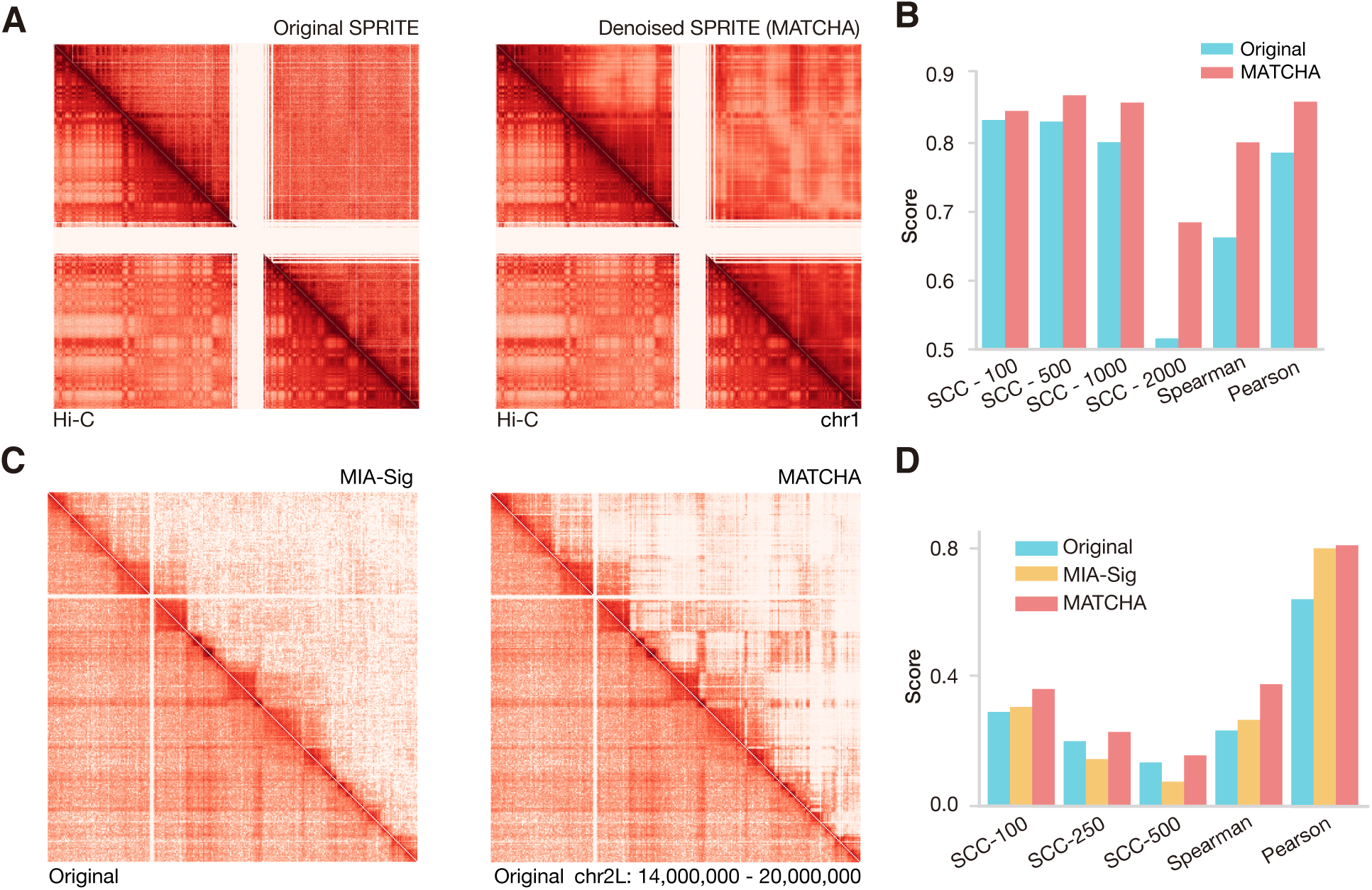
Evaluation of MATCHA’s performence in denoising multi-way chromatin interaction data. **(A)** Heatmap comparison of Hi-C versus the original SPRITE (left) and the denoised SPRITE by MATCHA (right). The heatmap is for chromosome 1 in GM12878 human cell line at 100kb resolution. **(B)** Similarity measurement of Hi-C versus the original SPRITE and the denoised SPRITE by MATCHA. The measurement includes the stratum adjusted correlation coefficient (SCC) with different maximum distance, Pearson correlation score, and Spearman correlation score. “SCC - *k*” stands for the SCC that considers the first *k* diagonals of the contact matrix. **(C)** Heatmap comparison of the original ChIA-Drop versus the ChIA-Drop denoised by MIA-Sig (left) and MATCHA (right). The heatmap is for chromosome 1 in *Drosophila* S2 cell line at 20kb resolution. **(D)** Similarity measurement of Hi-C versus the ChIA-Drop denoised by MIA-Sig and MATCHA. The measurement includes the SCC with different maximum distance, Pearson correlation score and Spearman correlation score. “SCC - *k*” stands for the SCC that considers the first *k* diagonals of the contact matrix.

Next, we generated the denoised contact map for the ChIA-Drop data at 20kb resolution. Here we used the ChIA-Drop data without ChIP enrichment (data from (Zheng et al., 2019)) to compare with the Hi-C data for *Drosophila* S2R+ (Szabo et al., 2018). Specifically, we compared the denoising results with a recently published ChIA-Drop denoising method, MIA-Sig (Kim et al., 2019). Fig. 3C shows the heatmap comparison of the original ChIA-Drop contact map versus the denoised contact maps by MIA-Sig and MATCHA, respectively. Similar to what we observed from the SPRITE data, as compared to the original contact map, both MATCHA and MIA-Sig make the chromatin interaction features clearer. In particular, for long-range interactions, both MIA-Sig and MATCHA remove more interactions as compared to the original ChIA-Drop data. However, the contact map produced by MATCHA yields even clearer patterns, especially for long-range contacts. We also used various metrics to assess the similarity between the contact map of Hi-C and the maps of denoised ChIA-Drop. As shown in Fig. 3D, MATCHA consistently achieves higher similarity scores with Hi-C as compared to the original contact map and the contact map produced by MIA-Sig. We also observed that, for the similarity scores that takes long-range interactions into account (SCC-250, SCC-500), MIA-Sig performs even worse than the original ChIA-Drop contact matrix, whereas MATCHA consistently outperforms the original ChIA-Drop contact matrix.

Taken together, these results further demonstrate that, by more reliably identifying multi-way chromatin interactions, MATCHA improves the overall data quality of SPRITE and ChIA-Drop.

### MATCHA can distinguish multi-way interactions from pairwise interaction cliques

The hyperedges defined from multi-way interaction data and the cliques defined from pairwise interaction data are drastically different. In particular, a hyperedge represents simultaneous interaction among chromatin loci in a single nucleus whereas a clique simply represents the presence of all pairwise interactions from a group of chromatin loci in a cell population. In other words, cliques do not have the single nucleus resolution. To test if MATCHA can effectively differentiate between hyperedges and cliques, we generated cliques based on Hi-C edges (referred to as Hi-C cliques) and required that the cliques satisfy the 1D genomic distance greater than 5Mb (same as the SPRITE data; see Methods). We provided these cliques to the trained model and evaluated the results by comparing to SPRITE. To prevent the model from “cheating” by memorizing all training data to achieve better performance, we reduced the number of training samples from 80% to 20% of the hyperedges, which were excluded in the following evaluation. After training the model, we predicted the probability score for the Hi-C cliques and grouped them based on the score. As shown in Fig. 4A, for Hi-C clique groups with higher probability scores, the percentage of cliques that are supported by SPRITE data is significantly higher than the groups with a lower score. Moreover, we also observed that the fraction of cliques that occur 3-7 times in SPRITE data, which were not included in the training data, are more enriched in groups with higher probability scores as well. This analysis suggests the ability of MATCHA to distinguish potential multi-way interactions from cliques based on population Hi-C data.

**Figure 4:**
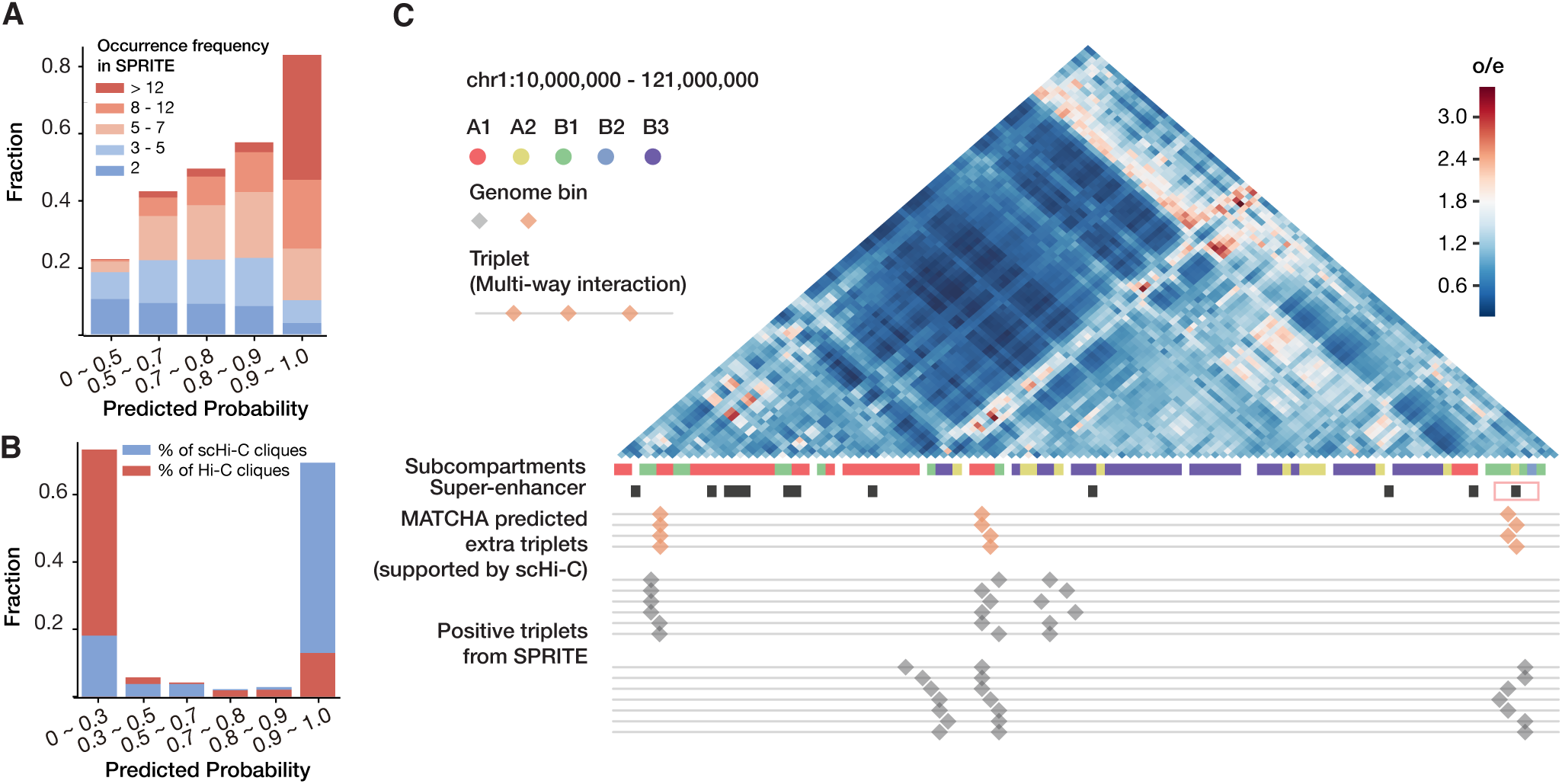
**(A)** Distribution of the occurrence frequency in the SPRITE data for the Hi-C cliques in each predicted probability group. **(B)** Overlapping between the Hi-C cliques with different predicted probability scores and the scHi-C cliques. The bar indicates the percentage of the total number of Hi-C cliques and scHi-C cliques in each group. **(C)** An example where the triplets predicted by MATCHA are not defined as positive samples based on occurrence frequency in SPRITE. The heatmap at the top shows the Hi-C O/E contact matrix. For the triplets track (bottom), each line represents a triplet either predicted by MATCHA or from the SPRITE data. Subcompartment and super-enhancer annotations for GM12878 are also shown. The relevant super-enhancer is marked in red rectangle. Note that there is a transcription factor GLIS1 expressed in the middle anchor region regulates 7 genes (ZBTB17, SPEN, FBLIM1, DDI2, AGMAT, CASP9, DNAJC16) in the left anchor region (not shown).

We further assessed if these Hi-C cliques with high probability scores indeed interact in the same nuclei by comparing with single cell Hi-C (scHi-C) data (Ramani et al., 2017). For the contact matrix of each single cell, we created an edge for each non-zero entry and then collected all cliques that satisfy the 1D genomic distance constraint (referred to as scHi-C cliques). These scHi-C cliques indicate that all pairwise interactions between these three bins happen in the same nucleus. In Fig. 4B, we found that of all the scHi-C cliques that overlap with the Hi-C cliques, about 70% of them are in the group with a probability score greater than 0.9, which makes up less than 20% of the Hi-C cliques. On the other hand, the Hi-C cliques with less than 0.3 probability scores account for more than 70% but only overlap with less than 20% of the scHi-C cliques. These results suggest that among all the Hi-C cliques, those received higher probability scores from MATCHA are enriched with multi-way interactions while the rest correspond to combinations of pairwise interactions within a cell population.

In Fig. 4C, we show an example of a Hi-C clique that receives a high probability score from MATCHA and overlaps with the scHi-C clique but was not defined as positive sample based on the occurrence frequency from SPRITE. The three regions predicted by MATCHA to form a triplet hyperedge are close to A1 and A2 active subcompartments (Rao et al., 2014). We refer to these three regions as the left, middle, and right anchor regions in Fig. 4C. Based on the super-enhancer annotations for GM12878 (Hnisz et al., 2013), one super-enhancer is at the right anchor region, which also belongs to the A2 subcompartment.

By comparing to the expressed genes in these three anchor regions and the transcriptional regulatory network (Marbach et al., 2016), we found that the transcription factor GLIS1 expressed in the middle anchor region regulates 7 genes in the left anchor region. Additional motif scan by FIMO (Grant et al., 2011) confirmed the existence of two GLIS1 binding motifs with *p*-value 1.34 × 10^−5^ and 1.02 × 10^−5^, respectively, within the super-enhancer at the right anchor region. Therefore, this potential three-way interaction among a super-enhancer, transcriptional factor, and target genes may reflect a higher-order module of chromatin interaction and transcriptional regulation (Stadhouders et al., 2019; Tian et al., 2020). We further analyzed the triplets from SPRITE with relatively low occurrence frequency (i.e., not positive samples) and found four each observed only twice that are close to this triplet (+/-2Mb for each anchor point), supporting the MATCHA prediction.

Collectively, these results demonstrate that the higher-order relationships in the multi-way chromatin interaction data are more than the combination of pairwise interactions and that MATCHA is able to model and predict multi-way contacts properly, leading to potential new insights into the interplay between transcription regulation and 3D genome organization.

### Embeddings produced by MATCHA reflect 3D genome function and spatial localization

To further demonstrate that MATCHA reliably extracts chromatin interaction patterns from the constructed hypergraph, we analyzed the embeddings produced by MATCHA trained from the SPRITE data. We first demonstrated the impact of the Mix-n-Match autoencoder (see Methods) by replacing it with the standard paired autoencoder. The model reached similar performance in predicting hyperedges, as expected. We then visualized the learned embeddings by projecting them to two-dimensional space with PCA. Each data point in Fig. 5A represents one 1Mb genomic bin with colors indicating the chromosome to which it belongs. We found that the embeddings of genomic bins based on the standard autoencoder form clusters according to the chromosome, making it impossible to make meaningful comparisons.

**Figure 5:**
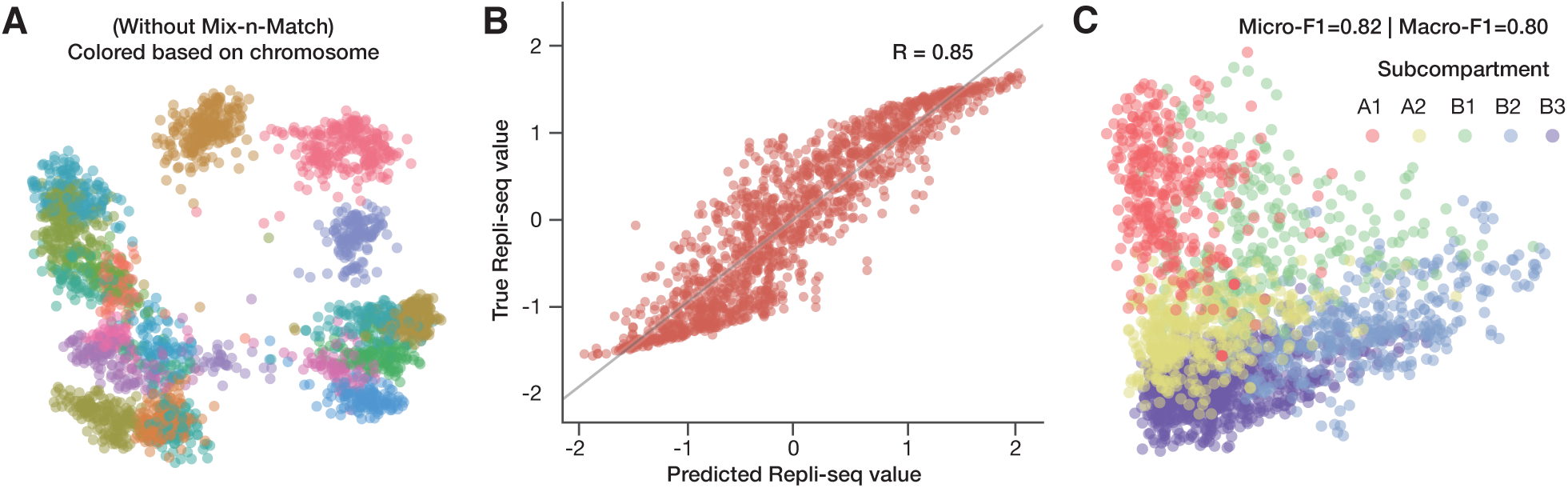
Evaluations for the learned embeddings of the genomic bins. **(A)** Visualization of the embeddings without the Mix-n-Match autoencoder. The embeddings are projected to two dimensions by PCA. Data points (genomic bins) are colored based on the chromosome they are from. Without the Mix-n-Match setting, genomic bins from the same chromosome are clustered. **(B)** Correlation of the predicted Repli-seq value using the learned embedding vectors versus the true Repli-seq value. **(C)** Visualization of the embeddings with the Mix-n-Match autoencoder. Data points are colored based on Hi-C subcompartment annotation.

We then evaluated the embeddings from MATCHA with the Mix-n-Match autoencoder by predicting the DNA replication timing based on Repli-seq signals. We binned the two-fraction Repli-seq signals to 1Mb resolution, and applied genome-wide z-score normalization. A linear regression model was trained with odd-numbered chromosomes and tested on the even-numbered chromosomes. The predicted value versus true signal value is shown in Fig. 5B, where a strong correlation can be observed (Pearson correlation = 0.85), suggesting that the MATCHA embeddings reflect the replication timing program, which is a fundamentally important genome function. Because the embeddings are enriched with the information extracted from the multi-way chromatin interaction data, even simple models like linear regression can reach a high correlation score. We tried more complicated regression models such as random forest regression and found the correlation score was similar to that achieved with linear regression.

We further asked if these embeddings capture Hi-C subcompartments, which are spatial genome segregation patterns in cell nucleus (Rao et al., 2014). Since the original subcompartment annotation is at 100Kb resolution, we converted it to the 1Mb resolution by a “voting scheme”. Specifically, for each 1Mb, there are 10 labels from 100Kb subcompartment annotations. When more than half of the labels belong to the same group, that 1Mb is labeled as the corresponding subcompartment. Otherwise, that bin would be removed in the next step. We also excluded the very small subcompartment B4 which only exists on chromosome 19. More than 95% of the bins received subcompartment labels. We again projected the embeddings to two-dimensional space using PCA and visualized the projected embedding vectors with annotations of subcomparments (Fig. 5C). We found that overall the genomic bins belonging to the same subcompartment are obviously clustered together. Since the subcompartment annotations reflect the spatial segregation pattern of the genome with a gradual change, as expected, there is no clear separation between clusters. This again demonstrates that by including the Mix-n-Match autoencoder, the bins are no longer clustered into groups merely based on the chromosome from which they come. Additionally, we quantitatively evaluated the consistency between embedding vectors with subcompartments by training a logistic regression to classify the genomic bins based on the embedding vectors. We used half of the genomic bins as training data and made sure that the bins from the same chromosome either appear all in the training data or all in the testing data. For the testing set, a simple logistic regression model can make accurate predictions with the micro-F1 (accuracy) score of 0.82 and the macro-F1 score of 0.80.

These results demonstrate that the Mix-n-Match autoencoder scheme in MATCHA is effective in forming genome-wide embeddings, which successfully capture genome structure and function based on multi-way interaction data.

## Discussion

Recent advances of ligation-free, genome-wide chromatin interaction mapping methods such as SPRITE (Quinodoz et al., 2018) and ChIA-Drop (Zheng et al., 2019) provide new perspectives on 3D genome and function by revealing multi-way contacts within the same nuclei (Kempfer and Pombo, 2019). However, computational methods that can fully utilize the potential of such multi-way chromatin interaction data remain underdeveloped. In this work, we developed MATCHA, a new multi-way chromatin interaction analysis framework based on hypergraph representation learning. We demonstrated that MATCHA can effectively extract multi-way chromatin interactions. Specifically, the method is able to make accurate predictions for multi-way interacting genomic loci to denoise the original data. We also showed the effectiveness of MATCHA by comparing with additional datasets such as Hi-C and ChIA-PET as well as its potential of identifying new multi-way interactions missed by the original data.

MATCHA has several algorithmic novelties: (1) To our knowledge, this is the first method that analyzes the multi-way chromatin interaction data based on hypergraph representation learning. MATCHA has strong promise in capturing the embeddings of multi-way interaction and can also be used to denoise input data. (2) We incorporated our recently developed Hyper-SAGNN (Zhang et al., 2020), a hypergraph representation learning paradigm into MATCHA. Specifically, we designed a novel feature generation method and a biologically-motivated negative sampling approach to make the model better suited for multi-way chromatin interaction. (3) We also enhanced the scalability of MATCHA with an efficient Bloom filter data structure that allows accurate and efficient negative sampling.

MATCHA can be further improved to better characterize higher-order interactions among different components in the nucleus. Although we mainly focused on analyzing the multi-way interaction among different genome loci (i.e. a homogeneous hypergraph as the nodes are all genomic bins), MATCHA can be extended to incorporate other constituents in the cell nucleus in addition to chromatin (e.g., proteins and RNAs) as a heterogeneous hypergraph based on emerging datasets such as RNA-DNA SPRITE (Quinodoz et al., 2018). Our hypergraph representation learning method Hyper-SAGNN can effectively learn the embeddings for heterogeneous hypergraphs (Zhang et al., 2020). In addition, we chose to train MATCHA using multi-way interaction data only in this work, but the model can easily include other functional genomic signals as features of the corresponding genomic bins. This would in principle extend the existing work on predicting pairwise chromatin loops based on functional genomic data (Huang et al., 2015; Whalen et al., 2016; Zhang et al., 2018; Kai et al., 2018; Zhang et al., 2019). Finally, the multi-way chromatin interactions extracted by MATCHA could also be the foundation to connect transcriptional regulation and 3D genome organization (Stadhouders et al., 2019; Kim and Shendure, 2019; Tian et al., 2020). Indeed, data from GAM (Beagrie et al., 2017) and SPRITE (Quinodoz et al., 2018) revealed the abundance of three-way interactions involving super-enhancer and active genes. Promoters may also act as enhancers to regulate other genes (Li et al., 2012; Engreitz et al., 2016). The recent data based on ChIA-Drop (Zheng et al., 2019) also uncovered three-way interactions among promoters that have imbalanced gene expression level in which the promoters with lower transcription level might act as enhancers. These observations further suggest the importance of investigating chromatin interactions in a non-pairwise manner. Taken together, we believe that MATCHA provides an effective algorithmic framework for the modeling and analysis of multi-way chromatin interaction data with the potential to advance our understanding of nuclear organization.

## Methods

### Definitions of hypergraph and the hyperedge prediction problem

#### Hypergraph

A hypergraph is defined as *G* = (*V, E*), where *V* = {*n*_1_, …, *n*_*n*_} represents the set of nodes in the graph, and 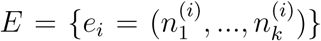 represents the set of hyperedges. For any hyperedge *e*, it connects two or more nodes (|*e*| ≥ 2). If all the hyperedges in a hypergraph contain the exact same number of nodes 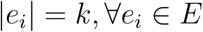, it is referred to as a *k*-uniform hypergraph.

#### The hyperedge prediction problem

The hyperedge prediction problem aims to learn a function *p* that can predict the probability of a group of nodes (*n*_1_, *n*_2_, …, *n*_*k*_) forming a hyperedge (Zhang et al., 2020):

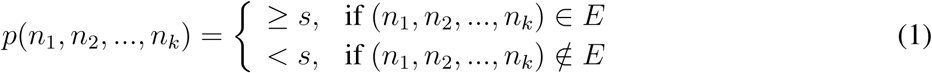

where *s*, typically chosen as 0.5, is the threshold to binarize the continuous probability score into a label indicating the existence of the corresponding hyperedge. As *n*_*i*_ is just the id for the corresponding node, to make the problem numerically tractable, it is natural to assume that the features of nodes 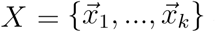 are known. This makes it possible to rewrite the function as:

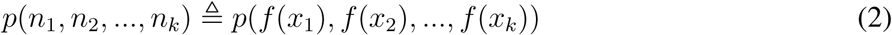

where the transformation of features *f* (*x*_*i*_) can be considered as the embedding vectors for the node *n*_*i*_.

### Data processing

The GM12878 DNA SPRITE cluster files on hg38 (Quinodoz et al., 2018) were downloaded from the 4DN data portal: https://data.4dnucleome.org. We also downloaded the processed RNAPII enriched ChIA-Drop data and ChIA-Drop data of *Drosophila* S2 cell from GSE109355 (Zheng et al., 2019). When creating the contact matrix for the SPRITE data, we used the same procedure in Quinodoz et al. (2018) for balancing the weight of each pairwise interaction with the size of the original SPRITE cluster. The contact matrix was further normalized by matrix balancing. For the ChIA-Drop data, we did not perform further normalization after decomposition.

The GM12878 *in-situ* Hi-C data on hg38 were downloaded from the 4DN data portal. We used KR matrix balancing for normalization. Two fraction Repli-seq data were also obtained from the 4DN data portal. The single cell Hi-C (scHiC) data of GM12878 were downloaded from GSE84920 (Ramani et al., 2017). For the scHi-C data, the aligned paired-end reads were converted to hg38 using UCSC liftOver. The alignment was then binned to produce the contact matrix for each single cell. To reduce the sparsity of the contact matrix, we applied linear convolution to each with a window size of 1 and step size of 1. The RNAPII enriched ChIA-PET data were downloaded from the same GSE repository as the ChIA-Drop data (Zheng et al., 2019). The processed *in-situ* Hi-C data for *Drosophila* S2 cell were downloaded from GSE99104 (Szabo et al., 2018).

### Constructing hypergraphs based on multi-way chromatin interaction data

For the SPRITE data, nodes in the graph represent non-overlapping 1Mb genomic bins (bins at the centromere are discarded). When certain genomic bins share multiple SPRITE clusters, we connect the corresponding nodes using a hyperedge. Note that we chose to focus on hyperedges with the size of 3 to 5 because they are more abundant in the dataset and can be filtered with high occurrence frequency to have enough number of samples for the model training. Specifically, we decomposed SPRITE clusters with size less than or equal to 25 into subsets of the corresponding size and counted the frequency of each combination of genomic bins (referred to as *k*-mer in the SPRITE paper (Quinodoz et al., 2018)). To reduce the number of *k*-mers to be considered, we focused on relatively distal interactions by requiring the minimum intra-chromosomal genomic distance *>*5Mb. We further used different occurrence frequency cut-off for different sized hyperedges to balance the average size of hyperedges. Note that in this work we did not include SPRITE clusters with size larger than 25 to reduce the processing time for decomposing clusters into hyperedges.

For the RNAPII ChIA-Drop data, a similar data processing procedure was applied with the resolution of 5Kb to generate hyperedges of size 3 to 5. Compared to SPRITE, *k*-mers in ChIA-Drop usually have smaller occurrence frequency; we therefore chose to use a uniform cut-off of 2 to define hyperedges and do not constrain the genomic distance. Since ChIA-Drop data only contains intra-chromosomal clusters, as a proof-of-principal, we evaluated our MATCHA method on ChIA-Drop data of chr2L and chr2R in the *Drosophila* genome.

### Labeled data generation

The Hyper-SAGNN model in MATCHA (see later section) was trained to be a binary classifier for the existence of the hyperedge among a group of nodes (Fig. 1A). The positive samples are the observed hyperedges in the constructed hypergraph while the negative samples are groups of nodes unobserved as the hyperedge. We can in principle generate random combinations of genomic bins and consider them as negative samples; however, this would oversimplify the prediction task because most of the randomly generated negative samples can be identified by simple metrics, e.g., whether it contains inter-chromosomal interactions or the 1D genomic distance. We therefore designed the following new procedure to generate negative samples by changing a small fraction of the nodes in the positive samples. For each positive sample of size *k*, the number of nodes to be altered *n* is sampled from a zero truncated binomial distribution with parameter *k* and a hyperparameter *p*, which is equivalent to altering each node in the positive samples independently with probability *p* while making sure at least one node would change, i.e.,

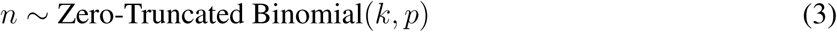

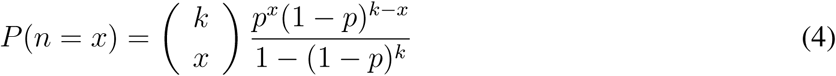

Smaller *p* would lead to smaller averaged difference between positive and negative samples, producing more difficult negative samples. Here we chose *p*=0.5 which leads to 1.7 to 2.5 expected difference between positive and negative samples for hyperedges of size 3 to 5. We then randomly selected *n* nodes and changed each of them. We required that the changed node is from the same chromosome to make the problem harder. We further made sure that this sample satisfies the genomic distance constraint. Specifically, for SPRITE, the minimum intra-chromosomal 1D distance within a group of nodes should be larger than 5 bins (same as the positive samples). For ChIA-Drop, we ensured that the changed node is within the ±20 bins to make the distance profile of positive and negative samples similar to each other. Note that we did not use the genomic distances as features in MATCHA, but this feature can be easily incorporated into our model to potentially further improve performance. Finally, we assessed if this modified sample happens to be the same as any other positive samples. If that happened, we repeated the above process until having a negative sample satisfying all conditions (see an efficient approach we implemented for this work in later section). For each batch of positive samples, we generated 3 times the amount of negative samples. The negative samples were generated dynamically for each training and evaluation epoch instead of being generated beforehand. This approach allows a more accurate characterization of the negative samples and prevents potential over-fitting.

### A new Mix-n-Match autoencoder for node features generation

Here we describe our approach to generating the features for the nodes in the hypergraph, i.e., 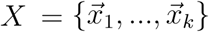. Although we can use functional genomic signals on the genomic bins (such as ChIP-seq for histone marks or transcription factors) as the node features, to demonstrate the generalization ability of this approach to cell types where these signals are inadequate, we developed a framework that generates features based on the hypergraph only (Fig. 1A).

We first decompose each hyperedge in the training data into pairwise interactions and create a corresponding adjacency matrix *A*. The *i*-th row of *A*, denoted by 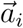, shows the neighborhood structures of the node *n*_*i*_, which then passes through an auto-encoder-like neural network to produce 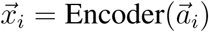. A decoder with symmetric structure is applied to reconstruct 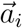 from 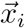. The corresponding mean-squared reconstruction error is added to the final loss term as a regularization term. The same strategy has been used in previous graph/hypergraph representation learning methods (Tu et al., 2018; Wang et al., 2016) and also in our recent work Hyper-SAGNN (Zhang et al., 2020), i.e,

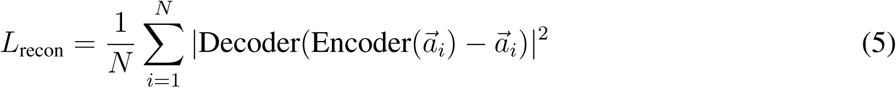

Note that, although this approach decomposes each hyperedge into pairwise interactions, the contact matrix is passed through the encoder which makes non-linear transformation of the input to be used as the node features for the predictions of higher order interactions. This significantly differs from the earlier work that decomposes the hyperedges into a contact matrix and studies the contact matrix directly. However, for networks constructed by chromatin interaction data, this encoder based approach does have a shortcoming. Data including Hi-C and SPRITE that have both intra-chromosomal and inter-chromosomal interactions usually contain more intra-chromosomal interactions. In other words, for each row in the genome-wide contact matrix, a small fraction of the columns (intra) receive more weight while a large fraction of the columns (inter) are much sparser and noisier. If we use the corresponding row in the genome-wide contact matrix as the feature, nodes from the same chromosome would have a more similar feature as compared to nodes from different chromosomes. Although this would not have a negative impact on hyperedge prediction, as nodes from different chromosomes indeed have very different spatial neighborhood structure in the graph, it may not be appropriate for analyzing the embedding vectors, as the model learned for one chromosome cannot generalize to the other ones.

We therefore designed a new method called “Mix-n-Match autoencoder”. A similar structure has been proposed in the computer vision field for image translation (Wang et al., 2018). For a genome with *n* chromosomes, we denote *C*_*ij*_ as the part of the genome-wide contact matrix corresponding to interactions between chromosome *i* and *j*. We use *n* encoder and *n* decoder where the *i*-th encoder Encoder_*i*_ takes vector of size *N*_*i*_ and produces a hidden vector of size *d*_*h*_, where *N*_*i*_ represents the number of bins in chromosome *i*. The Decoder_*i*_ works the other way around as the input size is *d*_*h*_. However, instead of making the encoder and decoder work in a paired manner (as described above), it is randomly paired (as the name ‘Mix-n-Match’ suggests). Specifically, for a node *k* from chromosome *i*, the *k*-th row in the intra-chromosomal contact matrix *C*_*ii*_ (denoted as 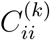) is taken as the input for Encoder_*i*_ to produce feature *x*_*k*_. Then a random chromosome *j, j* ≠ *i*, is selected with the corresponding decoder Decoder_*j*_ to reconstruct the *k*-th row in the inter-chromosomal contact matrix *C*_*ij*_ from the input *x*_*k*_. The new reconstruction loss for node *k* is therefore defined as:

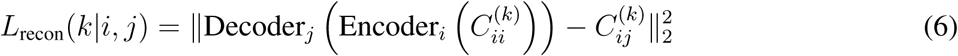

By adding this reconstruction loss to the final loss term, the model would make the embeddings for nodes from different chromosomes to be well “blended” as the same decoder *j* would be applied to nodes from all the other chromosomes to reconstruct inter-chromosomal interactions to chromosome *j*.

### The Hyper-SAGNN architecture for hypergraph representation learning

The detailed description of this part of the method can be found in our recent work Zhang et al. (2020). The structure of the neural network for Hyper-SAGNN is shown in Fig. 1B. The input to the model can be represented as tuples, i.e., 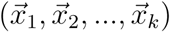. Each tuple first passes through a position-wise feed forward network to produce 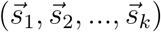, where 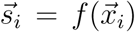. *f* represents the transformation of the neural network to 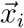. We refer to each 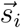 as the static embedding for node *i* since it remains the same for node *i* independent to the given tuple. The input also passes through another transformation to produce a new set of node embedding vectors 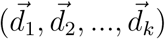, where 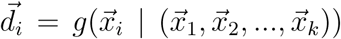. We refer to each 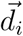 as the dynamic embedding because it depends on all the node features within this tuple. The transformation to produce dynamic embeddings is the multi-head self-attention layer (Vaswani et al., 2017) (see below):

Given a group of nodes 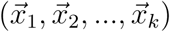 and weight matrices *W*_*Q*_, *W*_*K*_, *W*_*V*_ to be trained that represent the linear transformation of features before applying the scaled dot-product attention (Vaswani et al., 2017), the attention coefficients that indicate the pairwise importance of nodes are computed. These coefficients are then normalized through softmax to produce the final pairwise importance score:

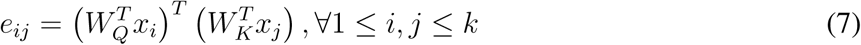

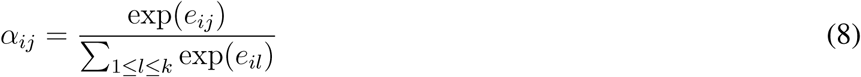

The dynamic embeddings are defined as the weighted sum of linear transformed features with a non-linear activation function:

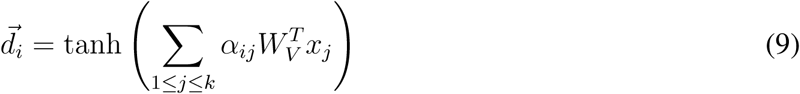

We further calculate the Hadamard power (element-wise power) of the difference of the corresponding static/dynamic pair for each node, which is subsequently passed through a one-layered neural network with sigmoid as the activation function to produce a probability score *p*_*i*_. Finally, all the output *p*_*i*_ ∈ [0, 1] are averaged to produce the final result *p*, i.e.,

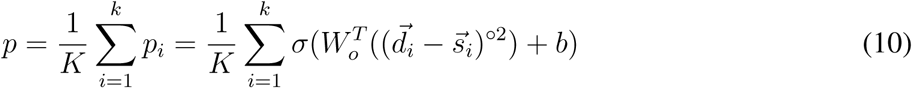

The Hyper-SAGNN model is trained to minimize the binary cross-entropy loss. The training procedure is terminated when it reaches a predefined number of training epochs or the performance stops improving on an individual validation set.

### Optimizing memory consumption via probabilistic data structure

One important practical issue in the negative sampling process (mentioned above) is that we ensure the generated negative samples do not overlap with known positive samples. Although one might argue that this process is unnecessary, as the probability for randomly generated negative samples being the same as a positive sample is small, this probability can increase greatly due to our stringent negative sampling strategy. Indeed, we found that the average number of trials for generating a non-overlapping negative sample is larger than 2.5 with the maximum trial number being more than 300. A similar technique has been developed in previous hypergraph representation learning methods (Tu et al., 2018) where a dictionary is maintained to keep the record of all positive samples. However, maintaining a dictionary in memory consumes enormous resources and significantly increases runtime for building it. This problem becomes even more significant when the number of hyperedges exceeds 2 million for SPRITE (as compared to 100K in the dataset that previous methods used (Tu et al., 2018)).

To reduce the memory consumption while maintaining an acceptable query runtime, we utilized the Bloom filter (Bloom, 1970) to keep track of the observed hyperedges. The Bloom filter is a type of data structure that returns whether an element is a member of a set and has several advantages. First, it is highly memory-efficient at the cost of producing potential false positives. However, this will have little impact for our purpose if we assume the false positive samples from the Bloom filter are distributed relatively evenly in the negative samples. Here, we control the error rate to be less than 10^−3^ by setting the number of hash functions and the size of the bit array. Second, it has constant runtime for the adding operation, which leads to a shorter wait time before the actual training process. In the extreme case, when it is not possible to maintain the Bloom filter in memory, we use the memory-mapped Bloom filter which allows us to keep the data structure in the hard disk and query (Debnath et al., 2011). Finally, the query process is still efficient as compared to searching algorithms over the training data.

In our implementation, before the labeled data generation step, we built a Bloom filter that stores all the positive samples and “potential” hyper-edges. For SPRITE, these are *k*-mers that have occurrence frequency greater than 2 but smaller than the chosen cut-off. For ChIA-Drop, these are *k*-mers that have occurrence frequency equal to 1. These samples cannot be classified into either positive or negative samples based on the current data, so these are excluded in the performance evaluation. By incorporating the Bloom filter into the implementation, our method is able to deal with large datasets or hyperedges of larger size efficiently. This would greatly enhance the scalability in practice.

## Acknowledgment

This work was supported in part by the National Institutes of Health Common Fund 4D Nucleome Program grant U54DK107965 (J.M.) and the National Institutes of Health grant R01HG007352 (J.M.). The authors would like to thank Ben Chidester, Minji Kim, and Yijun Ruan for helpful discussions.

## Author Contributions

Conceptualization, R.Z. and J.M.; Methodology, R.Z. and J.M.; Software, R.Z.; Investigation, R.Z. and J.M.; Writing – Original Draft, R.Z. and J.M.; Writing – Review & Editing, R.Z. and J.M.; Funding Acquisition, J.M.

## References

R. A. Beagrie, A. Scialdone, M. Schueler, D. C. Kraemer, M. Chotalia, S. Q. Xie, M. Barbieri, I. de Santiago, L.-M. Lavitas, M. R. Branco, et al. Complex multi-enhancer contacts captured by genome architecture mapping. Nature, 543(7646):519, 2017.

C. Berge. Hypergraphs: combinatorics of finite sets, volume 45. Elsevier, 1984.

B. H. Bloom. Space/time trade-offs in hash coding with allowable errors. Communications of the ACM, 13(7):422–426, 1970.

B. Bonev and G. Cavalli. Organization and function of the 3D genome. Nature Reviews Genetics, 17 (11):661, 2016.

B. Debnath, S. Sengupta, J. Li, D. J. Lilja, and D. H. Du. Bloomflash: Bloom filter on flash-based storage. In 2011 31st International Conference on Distributed Computing Systems, pages 635–644. IEEE, 2011.

J. R. Dixon, S. Selvaraj, F. Yue, A. Kim, Y. Li, Y. Shen, M. Hu, J. S. Liu, and B. Ren. Topological domains in mammalian genomes identified by analysis of chromatin interactions. Nature, 485(7398): 376, 2012.

J. M. Engreitz, J. E. Haines, E. M. Perez, G. Munson, J. Chen, M. Kane, P. E. McDonel, M. Guttman, and E. S. Lander. Local regulation of gene expression by lncrna promoters, transcription and splicing. Nature, 539(7629):452–455, 2016.

M. J. Fullwood and Y. Ruan. ChIP-based methods for the identification of long-range chromatin interactions. Journal of Cellular Biochemistry, 107(1):30–39, 2009.

C. E. Grant, T. L. Bailey, and W. S. Noble. Fimo: scanning for occurrences of a given motif. Bioinformatics, 27(7):1017–1018, 2011.

A. Grover and J. Leskovec. node2vec: Scalable feature learning for networks. In Proceedings of the 22nd ACM SIGKDD International Conference on Knowledge Discovery and Data Mining, pages 855–864. ACM, 2016.

H. Gui, J. Liu, F. Tao, M. Jiang, B. Norick, and J. Han. Large-scale embedding learning in heterogeneous event data. In 2016 IEEE 16th International Conference on Data Mining (ICDM), pages 907–912. IEEE, 2016.

D. Hnisz, B. J. Abraham, T. I. Lee, A. Lau, V. Saint-André, A. A. Sigova, H. A. Hoke, and R. A. Young. Super-enhancers in the control of cell identity and disease. Cell, 155(4):934–947, 2013.

J. Huang, E. Marco, L. Pinello, and G.-C. Yuan. Predicting chromatin organization using histone marks. Genome Biology, 16(1):162, 2015.

Y. Kai, J. Andricovich, Z. Zeng, J. Zhu, A. Tzatsos, and W. Peng. Predicting CTCF-mediated chromatin interactions by integrating genomic and epigenomic features. Nature Communications, 9(1):4221, 2018.

R. Kempfer and A. Pombo. Methods for mapping 3D chromosome architecture. Nature Reviews Genetics, doi: 10.1038/s41576-019-0195-2, 2019.

M. Kim, M. Zheng, S. Z. Tian, B. Lee, J. H. Chuang, and Y. Ruan. MIA-Sig: multiplex chromatin interaction analysis by signal processing and statistical algorithms. Genome Biology, 20(1):251, 2019.

S. Kim and J. Shendure. Mechanisms of interplay between transcription factors and the 3D genome. Molecular Cell, 2019.

R. I. Kumaran, R. Thakar, and D. L. Spector. Chromatin dynamics and gene positioning. Cell, 132(6): 929–934, 2008.

G. Li, X. Ruan, R. K. Auerbach, K. S. Sandhu, M. Zheng, P. Wang, H. M. Poh, Y. Goh, J. Lim, J. Zhang, et al. Extensive promoter-centered chromatin interactions provide a topological basis for transcription regulation. Cell, 148(1-2):84–98, 2012.

E. Lieberman-Aiden, N. L. Van Berkum, L. Williams, M. Imakaev, T. Ragoczy, A. Telling, I. Amit, B. R. Lajoie, P. J. Sabo, M. O. Dorschner, et al. Comprehensive mapping of long-range interactions reveals folding principles of the human genome. Science, 326(5950):289–293, 2009.

D. Marbach, D. Lamparter, G. Quon, M. Kellis, Z. Kutalik, and S. Bergmann. Tissue-specific regulatory circuits reveal variable modular perturbations across complex diseases. Nature Methods, 2016.

E. P. Nora, B. R. Lajoie, E. G. Schulz, L. Giorgetti, I. Okamoto, N. Servant, T. Piolot, N. L. van Berkum, J. Meisig, J. Sedat, et al. Spatial partitioning of the regulatory landscape of the X-inactivation centre. Nature, 485(7398):381, 2012.

B. Perozzi, R. Al-Rfou, and S. Skiena. Deepwalk: Online learning of social representations. In Proceedings of the 20th ACM SIGKDD international conference on Knowledge Discovery and Data Mining, pages 701–710. ACM, 2014.

S. A. Quinodoz, N. Ollikainen, B. Tabak, A. Palla, J. M. Schmidt, E. Detmar, M. M. Lai, A. A. Shishkin, P. Bhat, Y. Takei, et al. Higher-order inter-chromosomal hubs shape 3D genome organization in the nucleus. Cell, 174(3):744–757, 2018.

V. Ramani, X. Deng, R. Qiu, K. L. Gunderson, F. J. Steemers, C. M. Disteche, W. S. Noble, Z. Duan, and J. Shendure. Massively multiplex single-cell Hi-C. Nature Methods, 14(3):263, 2017.

S. S. Rao, M. H. Huntley, N. C. Durand, E. K. Stamenova, I. D. Bochkov, J. T. Robinson, A. L. Sanborn, I. Machol, A. D. Omer, E. S. Lander, et al. A 3D map of the human genome at kilobase resolution reveals principles of chromatin looping. Cell, 159(7):1665–1680, 2014.

T. J. Salameh, X. Wang, F. Song, B. Zhang, S. M. Wright, C. Khunsriraksakul, and F. Yue. A supervised learning framework for chromatin loop detection in genome-wide contact maps. bioRxiv, page 739698, 2019.

R. Stadhouders, G. J. Filion, and T. Graf. Transcription factors and 3D genome conformation in cell-fate decisions. Nature, 569(7756):345–354, 2019.

Q. Szabo, D. Jost, J.-M. Chang, D. I. Cattoni, G. L. Papadopoulos, B. Bonev, T. Sexton, J. Gurgo, C. Jacquier, M. Nollmann, et al. TADs are 3D structural units of higher-order chromosome organization in Drosophila. Science Advances, 4(2):eaar8082, 2018.

Z. Tang, O. J. Luo, X. Li, M. Zheng, J. J. Zhu, P. Szalaj, P. Trzaskoma, A. Magalska, J. Wlodarczyk, B. Ruszczycki, et al. CTCF-mediated human 3D genome architecture reveals chromatin topology for transcription. Cell, 163(7):1611–1627, 2015.

D. Tian, R. Zhang, Y. Zhang, X. Zhu, and J. Ma. MOCHI enables discovery of heterogeneous interactome modules in 3D nucleome. Genome Research, pages gr–250316, 2020.

K. Tu, P. Cui, X. Wang, F. Wang, and W. Zhu. Structural deep embedding for hyper-networks. In Thirty-Second AAAI Conference on Artificial Intelligence, 2018.

A. Vaswani, N. Shazeer, N. Parmar, J. Uszkoreit, L. Jones, A. N. Gomez, Ł. Kaiser, and I. Polosukhin. Attention is all you need. In Advances in Neural Information Processing Systems, pages 5998–6008, 2017.

D. Wang, P. Cui, and W. Zhu. Structural deep network embedding. In Proceedings of the 22nd ACM SIGKDD International Conference on Knowledge Discovery and Data Mining, pages 1225–1234. ACM, 2016.

Y. Wang, J. van de Weijer, and L. Herranz. Mix and match networks: encoder-decoder alignment for zero-pair image translation. In Proceedings of the IEEE Conference on Computer Vision and Pattern Recognition, pages 5467–5476, 2018.

S. Whalen, R. M. Truty, and K. S. Pollard. Enhancer–promoter interactions are encoded by complex genomic signatures on looping chromatin. Nature Genetics, 48(5):488, 2016.

K. Xiong and J. Ma. Revealing Hi-C subcompartments by imputing inter-chromosomal chromatin interactions. Nature Communications, 10, 2019.

T. Yang, F. Zhang, G. G. Yardimci, F. Song, R. C. Hardison, W. S. Noble, F. Yue, and Q. Li. HiCRep: assessing the reproducibility of Hi-C data using a stratum-adjusted correlation coefficient. Genome Research, 27(11):1939–1949, 2017.

R. Zhang, Y. Wang, Y. Yang, Y. Zhang, and J. Ma. Predicting CTCF-mediated chromatin loops using CTCF-MP. Bioinformatics, 34(13):i133–i141, 2018.

R. Zhang, Y. Zou, and J. Ma. Hyper-SAGNN: a self-attention based graph neural network for hypergraphs. In International Conference on Learning Representations (ICLR), 2020.

S. Zhang, D. Chasman, S. Knaack, and S. Roy. In silico prediction of high-resolution Hi-C interaction matrices. Nature Communications, 10(1):1–18, 2019.

M. Zheng, S. Z. Tian, D. Capurso, M. Kim, R. Maurya, B. Lee, E. Piecuch, L. Gong, J. J. Zhu, Z. Li, et al. Multiplex chromatin interactions with single-molecule precision. Nature, 566(7745):558, 2019.

D. Zhou, J. Huang, and B. Schölkopf. Learning with hypergraphs: Clustering, classification, and embedding. In Advances in Neural Information Processing Systems, pages 1601–1608, 2007.

